# In amygdala we trust: different contributions of the basolateral and central amygdala in learning whom to trust

**DOI:** 10.1101/2021.05.03.442429

**Authors:** Ronald Sladky, Federica Riva, Lisa Rosenberger, Jack van Honk, Claus Lamm

## Abstract

Human societies are built on cooperation and mutual trust, but not everybody is trustworthy. Research on rodents suggests an essential role of the basolateral amygdala (BLA) in learning from social experiences (Hernandez-Lallement J et al., 2016), which was also confirmed in human subjects with selective bilateral BLA damage as they failed to adapt their trust behavior towards trustworthy vs. untrustworthy interaction partners (Rosenberger LA et al., 2019). However, neuroimaging in neurotypical populations did not consistently report involvement of the amygdala in trust behavior. This might be explained by the difficulty of differentiating between amygdala’s structurally and functionally different subnuclei, i.e., the BLA and central amygdala (CeA), which have even antagonistic features particularly in trust behavior (van Honk J et al., 2013). Here, we used fMRI of the amygdala subnuclei of neurotypical adults (n=31f/31m) engaging in the repeated trust game. Our data show that both the BLA and the CeA play a role and indeed differentially: While the BLA was most active when obtaining feedback on whether invested trust had been reciprocated or not, the CeA was most active when subjects were preparing their next trust decision. In the latter phase, improved learning was associated with higher activation differences in response to untrustworthy vs. trustworthy trustees, in both BLA and CeA. Our data not only translate to rodent models and support our earlier findings in BLA-damaged subjects, but also show the specific contributions of other brain structures in the amygdala-centered network in learning whom to trust, and better not to trust.

**SIGNIFICANCE STATEMENT:** In this fMRI study, the central amygdala was found active during trust behavior planning, while the basolateral amygdala was active during outcome evaluation. When planning trust behavior, central and basolateral amygdala activation differences between the players was related to whether participants learned to differentiate the players’ trustworthiness. Nucleus accumbens tracked whether trust was reciprocated but was not related to learning. This suggests learning whom to trust is not related to reward processing in the nucleus accumbens but rather to engagement of the basolateral amygdala. This study overcomes major empirical gaps between animal models and human neuroimaging and shows how different amygdala subnuclei and connected areas orchestrate learning to form different subjective trustworthiness beliefs about others and guide trust choice behavior.

## INTRODUCTION

Human societies are built on cooperation and mutual trust. On the individual level, trusting another person entails potential rewards, but also risks if the other person is abusing our trust to our own disadvantage. Thus, learning to distinguish the trustworthiness of an interaction partner is important for successful social interactions. Research on rodents suggests an essential role of the basolateral amygdala (BLA) in learning from social experiences (Hernandez-Lallement J,van Wingerden M,Schäble S and Kalenscher T, 2016). In line with this, we showed in a previous study that human participants with selective bilateral BLA damage failed to adapt their trust behavior towards trustworthy vs. untrustworthy interaction partners in a repeated trust game (Rosenberger LA,Eisenegger C,Naef M,Terburg D,Fourie J,Stein DJ and van Honk J, 2019). However, functional reorganization after developmental brain damage might confine the generalizability of these findings to neurotypical populations. Neuroimaging in neurotypical populations indeed did not consistently report involvement of the amygdala in trust behavior. This might be explained by difficulties in differentiating between the amygdala’s structurally and functionally different subnuclei, i.e., the BLA and central amygdala (CeA), which have even antagonistic features particularly in trust behavior (van Honk J,Eisenegger C,Terburg D,Stein DJ and Morgan B, 2013).

The amygdala is widely regarded as paramount for social cognition (Adolphs R, 2010), but it has been investigated as a uniform structure in the majority of human neuroimaging studies (Gupta R et al., 2011). While this approach may be due to the limited spatial specificity of functional MRI particularly in the ventral brain (Sladky R et al., 2013;Sladky R et al., 2018), it ignores the structural and functional heterogeneity of this brain area and its subnuclei (Balleine BW and Killcross S, 2006). Here, we overcame the limitations of previous research by using an acquisition protocol optimized for imaging ventral brain areas (Robinson S et al., 2004) in combination with a multiband EPI sequence with high spatial and temporal resolution (Moeller S et al., 2010), allowing for a time-resolved analysis of amygdalar subnuclei.

Our recent research in participants with basolateral amygdala lesions (Rosenberger LA,Eisenegger C,Naef M,Terburg D,Fourie J,Stein DJ and van Honk J, 2019) proposed that a network centered around the basolateral amygdala adaptively subserves learning to trust and distrust others. Importantly, this novel insight was based on a trust game task in which the participants repeatedly interacted with a trustworthy and an untrustworthy interaction partner. The task thus allowed us to investigate the dynamics of trust formation, as well as the role that different decision-making processes play in that. Here, using functional MRI in a healthy neurotypical population we employ the exact same behavioral paradigm to confirm and extend these findings to the specific functions of the separate subnuclei of the amygdala and the networks they are a part of. Our main aims were to derive what role the different subnuclei of the amygdala play for different aspects relevant in learning whom to trust, and to link them to neural activation in other sub-cortical regions that are highly connected with the amygdala (i.e., the bed nucleus of the stria terminalis, the nucleus accumbens, and the substantia nigra/VTA) (Janak PH and Tye KM, 2015).

## MATERIALS AND METHODS

### PARTICIPANTS

62 heathy, neurotypical volunteers (age=23.83±3.15 years, f/m=31/31), mostly undergraduate students from Vienna, Austria were recruited. Exclusion criteria were standard MRI exclusion criteria (e.g.: pregnancy, claustrophobia, and MRI-incompatible implants, clinically significant somatic diseases), a history of psychiatric or neurological disorders, substance abuse, psychopharmacological medication, less than nine years of education, as well as not being task-naive (e.g., having already participated in a similar study or being a psychology student). All participants provided written informed consent in accordance with the Declaration of Helsinki and were compensated for their participation. The study was approved by the ethics committee of the Medical University of Vienna (EK-Nr. 1489/2015).

### PROCEDURE AND TASK

This study was part of a bigger project including two additional tasks and a sample of older adults, which are not reported in the current article. Participants were first invited to a screening session where they performed some cognitive tasks and filled in some self-reported measures of psychological traits. The main session was usually conducted within two weeks from the screening session. Participants were welcomed to the MRI facility (University of Vienna MR Center) together with two other participants, who were in fact two confederates of the experimenter invited to play the trustees’ role. After having signed the consent form and filled in the MR safety questionnaire, participants and confederates were introduced to the protocol of the whole session. Afterwards, they went through the training of the three tasks, including the trust game. At the end of the training, participants were required to answer some questions in order to make sure they understood the task. Participants were finally placed into the MR scanner, while the confederates were putatively playing the task in the computer room next to the scanner room.

The repeated trust game was adapted from our previous study (Rosenberger LA,Eisenegger C,Naef M,Terburg D,Fourie J,Stein DJ and van Honk J, 2019) and programmed in z-Tree (version 3.3.7; (Fischbacher U, 2007)). The script of this trust game is deposited online (Rosenberger LA,Eisenegger C,Naef M,Terburg D,Fourie J,Stein DJ and van Honk J, 2019). In short, two players per round, an investor and a trustee, exchange monetary units with the aim to maximize their monetary outcome. In total, 40 rounds were played and the participant always played the role of the investor, while the trustees were allegedly played by the two confederates in an alternate randomized order. In reality, the actions taken by the two trustees were preprogrammed in a way that one of the confederates was behaving in a trustworthy and the other one in an untrustworthy way. Confederates/trustees were of similar age and same gender as the participant. At the beginning of each round (i.e., 20 per trustworthy condition and 20 per untrustworthy condition) both players received an endowment of 10 monetary units. Then each round encompasses four phases. In the *preparation* phase, participants are presented with the picture of the trustee’s face they are playing with in the current round. In the *investment* phase, participants invest (part of) their endowment (at least 1 unit) and the investment is tripled and then transferred to the trustee. During the *waiting* phase, the trustees ostensibly perform their back-transfers. Finally, during the *outcome* phase, participants are presented with the back-transfer outcome. In the first two rounds, both the trustworthy and untrustworthy trustees back-transferred the same amount of the money invested to the participants. In the following rounds, the trustworthy trustee always back-transferred as much or more than the money invested by the player, whereas the untrustworthy trustee always back-transferred less than or as much as the money invested by the investor. The sums invested by the participants were considered as a measure of trust given to the two trustees by the participants and used as the main variable of interest. Points earned throughout the task were transformed to Euros and added to the participants’ compensation.

At the end of the task, participants were presented with the trustees’ picture and were asked to rate them on four adjectives: trustworthiness, fairness, attractiveness, and intelligence (original German: *Wie vertrauenswürdig/attraktiv/intelligent/fair haben Sie den/die Teilnehmer/in wahrgenommen?*). Ratings were provided on visual analogue scales and transformed off-line to a numerical range between −10 and +10.

### FUNCTIONAL MRI DATA ACQUISITION AND PROCESSING

MRI acquisitions were performed on a Skyra 3 Tesla MRI scanner (Siemens Healthineers, Erlangen, Germany) using the manufacturer’s 32 channel head coil at the MR Center of the University of Vienna. In a single session, one run of the repeated trust game was performed by the participant while we performed functional MRI using a gradient echo T2*-weighted echo planar image sequence with the following parameters: MB-EPI factor=4, TR/TE = 704/34 ms, 2.2×2.2×3.5 mm^3^, 96×92×32 voxels, flip angle=50°, n<2400 volumes.

Data processing and analyses of the functional MRI data were performed in SPM (SPM12, http://www.fil.ion.ucl.ac.uk/spm/software/spm12/) and the Python projects nipype (http://nipy.org/nipype) and nilearn (http://nilearn.github.io). Preprocessing comprised slice-timing correction (Sladky R et al., 2011), realignment, non-linear normalization of the EPI images to MNI space (final resolution = 1.5×1.5×1.5mm^3^) using ANTs (Avants BB et al., 2011), and spatial smoothing with a 6mm FWHM Gaussian kernel.

## EXPERIMENTAL DESIGN AND STATISTICAL ANALYSES

### Behavioral data analysis

It is commonly understood that participants’ investment behavior is a behavioral expression of how they judged the trustees’ trustworthiness and changes reflect the extent to which they updated their beliefs (Bellucci G et al., 2017;Chang LJ et al., 2010;Rosenberger LA,Eisenegger C,Naef M,Terburg D,Fourie J,Stein DJ and van Honk J, 2019). This *objective* measure of trust was used to distinguish between learners and non-learners (using the median as cut-off value) and for a Spearman correlation analysis between the subjective ratings (trustworthiness, fairness, attractiveness, and intelligence) and the BOLD response in the amygdala.

### Functional MRI data analysis

First-level analyses of the data were implemented using nipype and performed using SPM12’s GLM approach. The GLM design matrix encompassed individual regressors for each of the 4 task phases (i.e., preparation, investment, waiting, and outcome) and each of the 2 interaction partners (trustworthy and untrustworthy, resulting in 8 effects of interest. Additionally, 6 realignment parameters were added as nuisance regressors to account for residual head motion effects. Second-level analyses of the data were implemented using nipype and performed using SPM12’s group-level approach for visual inspection of the whole brain results.

Volume of interest analyses were performed on the mean timeseries extracted using nilearn’s fit_transform from anatomical masks from the BLA, CeA (Tyszka JM and Pauli WM, 2016), NAc (AAL Atlas), BNST (Torrisi S et al., 2015), and SN/VTA (Talairach atlas transformed to MNI space). To investigate phase-dependent activation, timeseries analyses were conducted using custom python scripts that reproduced SPM’s default GLM analysis, using SPM’s canonical HRF to convolve the regressors and a high-pass filter with the default f=1/128 Hz cut-off frequency to account for signal drifts. Comparisons between learners and non-learners were performed using two-sampled *t*-tests and based on their Spearman correlations.

To verify that sensitivity of the fMRI dataset was sufficient to distinguish between BLA and CeA activation, a functional connectivity analysis was conducted. Task fMRI data were corrected for white matter and CSF signal and task effects (Ganger S et al., 2015) using regression before estimation of the functional connectivity maps of the BLA and CeA seeds.

## RESULTS

Participants played the repeated trust game inside the MRI scanner with a trustworthy and an untrustworthy trustee, both simulated (2×20 rounds). In general, participants were able to adapt their trust behavior, i.e., investments in the trust game, to the trustworthy and the untrustworthy trustee. However, there was a marked variability within our study sample, which allowed for a partition into a *learner* and *non-learner* sub-group (**FIGURE 1**). The task consisted of four different task phases (i.e., the *preparation, investment, waiting*, and *outcome* phase). A detailed time-resolved analysis of the BLA and CeA revealed that activation changed over the course of the different task phases. We found maximum BLA activation in the *outcome* evaluation phase and maximum CeA activation in the *preparation* phase. Yet, there was no overall BLA and CeA activation difference between the trustworthy or untrustworthy trustee in any of the task phases (**FIGURE 2**). However, when differentiating between learners and non-learners, we observed more activation in the BLA and the CeA for the untrustworthy trustee during the *introduction* phase of a trust game round (**FIGURE 3**). Additionally, while nucleus accumbens (NAc), substantia nigra and ventral tegmental area (SN/VTA), and bed nucleus of the stria terminalis (BST) activity was increased for the trustworthy trustee during *outcome* evaluation, there was no group difference between learners and non-learners (**FIGURE 4**).

**FIGURE 1.**
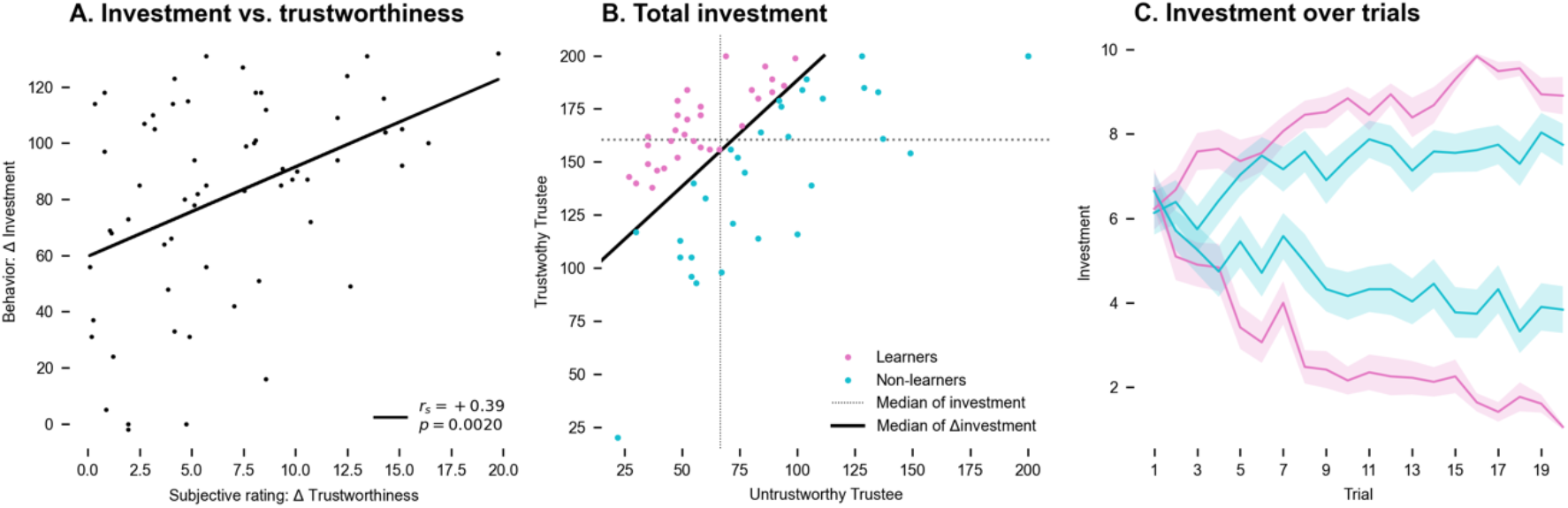
A. Investment vs. trustworthiness. Behavioral trust (***Δ*** investment) correlates with subjective ratings (***Δ*** trustworthiness rating), *r*_*s*_=+0.39, *p*=0.002. **B. Participants’ investment behavior**. In total, participants invested more in the trustworthy trustee. The difference between the investment into the trustworthy and untrustworthy trustee (***Δ*** investment) was used to median-split the population into a subgroup that learned to differentiate (learners, magenta color) and those who did not (non-learners, cyan color). **C. Participants’ investment behavior over time**. After a few trials, learners adapted their investment behavior to favor the trustworthy trustee. This differentiation was reduced in non-learners. Plot displays mean and SEM.

**FIGURE 2.**
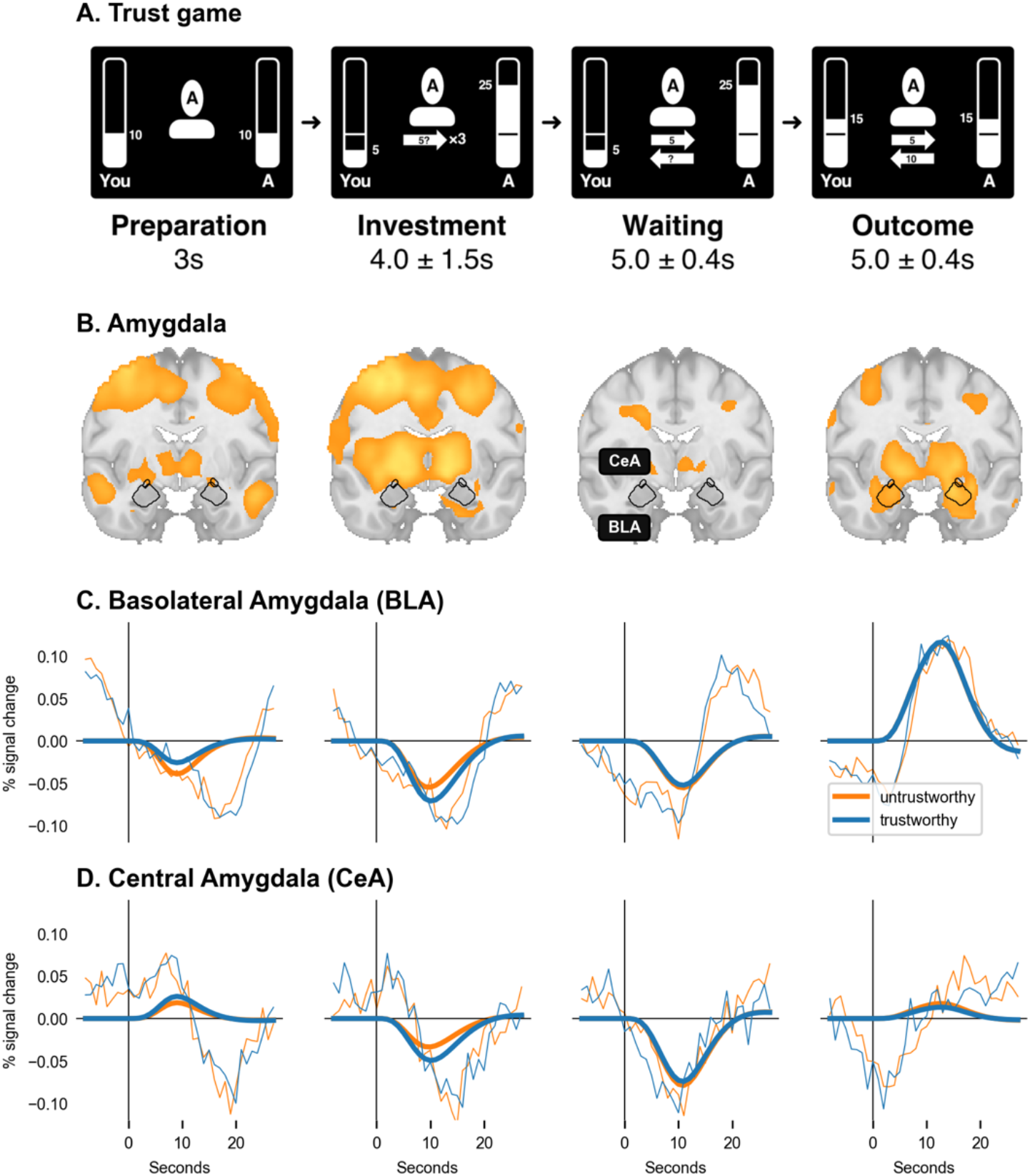
A. fMRI implementation of the trust game. Inside the MRI scanner, participants played the repeated trust game alternating with a (simulated) trustworthy and an untrustworthy trustee (2×20 rounds). *Preparation Phase*. Participants were presented with the face of the trustee they played with in this round. Both received an endowment of 10 points at the outset of each round. *Investment Phase*. Participants were asked to select an amount of 1 to 10 points to invest in the present trustee. The amount invested was tripled and added to the trustee’s account. *Waiting Phase*. While the trustees made their decision, the participant needed to wait. *Outcome Phase*. Finally, the trustee transferred back points to the participant, resulting in a non-negative outcome for the trustworthy (as shown in the example) and a non-positive outcome for the untrustworthy trustee. **B. Statistical parametric maps (SPMs) and outline of the anatomically defined Volumes of Interest (VOIs) of BLA and CeA**. SPMs show contrast for both trustees combined vs. baseline and are thresholded at p<0.001 for display purposes. **C & D. Time course of BLA and CeA BOLD responses**. CeA but not BLA was activated during the *preparation* phase, while BLA but not CeA was activated during the *outcome* phase. There were no activation differences between the trustworthy trustee (blue) and the untrustworthy trustee (orange). Thick lines represent the estimated BOLD model and fine lines represent the actual data (average VOI time courses).

**FIGURE 3.**
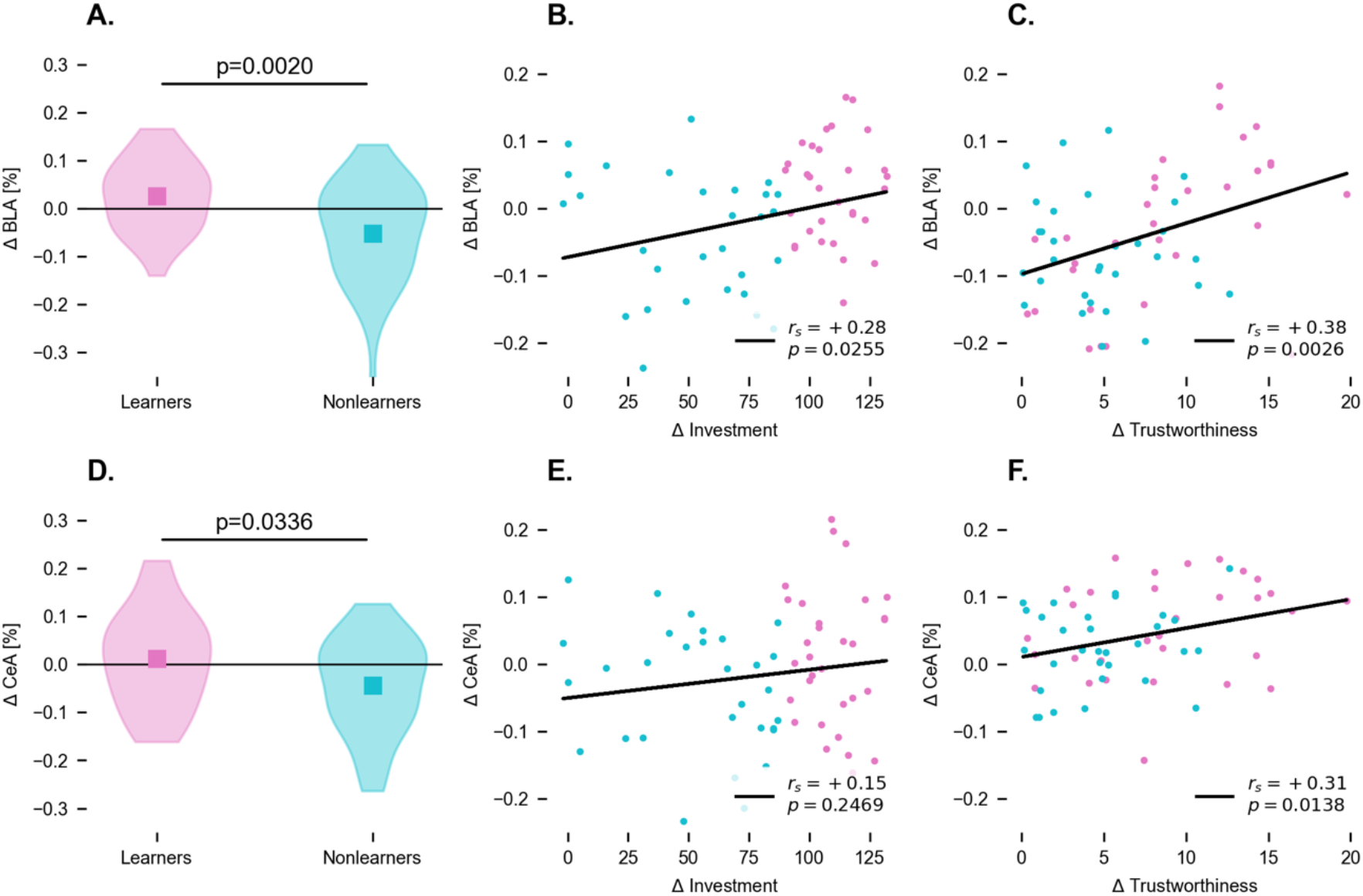
Activation differences between untrustworthy and trustworthy trustee in the *preparation* phase. BLA activation differences (contrast: untrustworthy - trustworthy) were higher for learners (magenta) vs. non-learners (cyan) (**A**), correlated with investment differences (**B**) and post-experiment subjective trustworthiness rating differences (**C**). The same relationship was found for CeA (**D & F**), except the correlation with investment differences was not significant (**E**).

**FIGURE 4.**
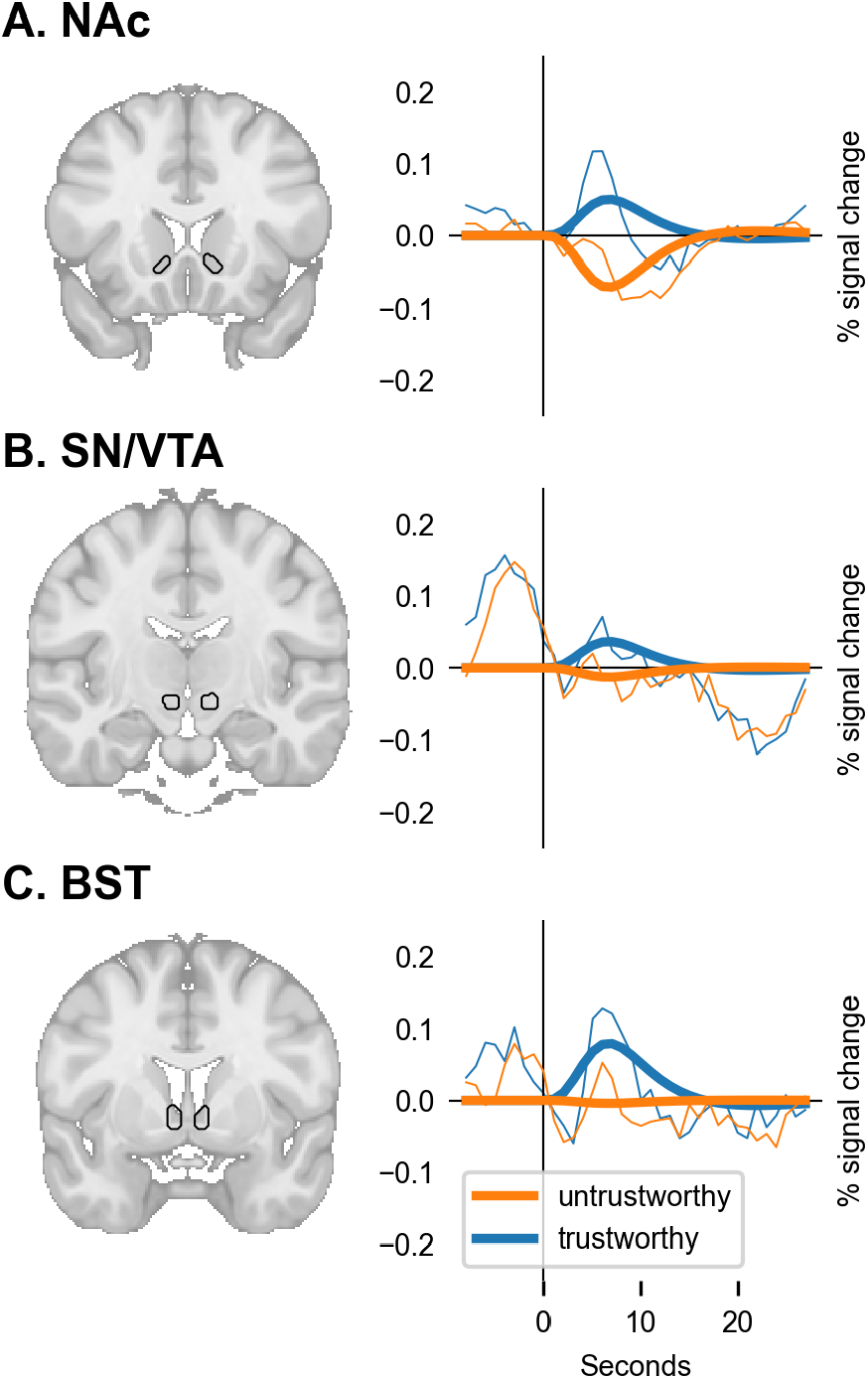
More activity for the trustworthy (blue) vs. untrustworthy trustee (orange) during the *outcome* event. (**A**) in the nucleus accumbens (NAc), T=+7.21, p<0.0001, (**B**) the substantia nigra (SN) and ventral tegmental area (VTA), T=+3.31, p<0.0010, and (**C**) the bed nucleus of the stria terminalis (BST), T=+4.38, p<0.0001. Thick lines represent the estimated BOLD model.

### BEHAVIORAL RESULTS

Marked trust differences emerged across the whole sample in the investment behavior towards the trustworthy as opposed to the untrustworthy trustee, with participants generally investing more in the trustworthy trustee on average, and increasingly so over the course of the repeated rounds of the task (**FIGURE 1B & C**). Morever, we find that individual differences in behavioral trust(*Δ investment* = *investment*_trustworthy_ - *investment*_untrustworthy_) showed a positive correlation with subjective trustworthiness ratings (*Δ trustworthiness* = *trustworthiness*_trustworthy_ - *trustworthiness*_untrustworthy_), *r*_*s*_ = +0.39, *p*=0.002 (**FIGURE 1A**). On the subjective level, the trustworthy trustee was rated as significantly more trustworthy, fair, and intelligent than the untrustworthy trustee (all p<0.05, Bonferroni corrected), but not as more attractive (n.s., after Bonferroni correction).

### NEUROIMAGING RESULTS

We find that different subnuclei of the amygdala were engaged in the trust game show increased activation during different phases of the task paradigm. This suggests that they are supposedly related to different aspects and processes required by the formation of trust. The two subnuclei that played the most specific role (**FIGURE 2B**) were the basolateral (BLA) and the central amygdala (CeA). Notably, the activation differences in these subnuclei and the validity of our analysis approach is supported by differences in their functional connectivity profiles, determined in our data. While the BLA connected to sensory integration areas and lateral PFC, the CeA connected to the ventral striatum, including the nucleus accumbens, and areas in the medial PFC (**SUPPLEMENTARY FIGURE 1**). The role of these subnuclei in the different task phases is as follows.

In the *preparation* phase, activity in the BLA was reduced (*T*=−8.9, *p*<0.0001) and in the CeA increased (*T*=9.9, *p<*0.0001), compared to the fixation baseline. Both BLA and CeA activity were reduced during *investment* (*T*=−15.4, *p*<0.0001 and *T*=−9.5, *p*<0.0001) and *waiting* phase (*T*=−9.7, *p*<0.0001 and *T*=−13.0, *p*<0.0001). During the *outcome* evaluation phase, activity in the BLA was increased (*T*=14.3, *p*<0.0001), while it was reduced in the CeA (*T*=−3.1, *p*=0.002232) (**FIGURE 2C & 2D**). All reported *p*-values survive Bonferroni correction for multiple comparisons (at p<0.05 Bonferroni FWE-corrected). Note that these response patterns were irrespective of whether a participant played with a trustworthy or untrustworthy trustee, as there were no significant differences between these two conditions. These findings thus relate to the general role of the amygdala subnuclei in the different parts of the task, and the overall processes and subfunctions engaged by the trust decision.

As a next step, we aimed to pinpoint how the engagement of the amygdala was related to differential evaluations of trustworthiness, and the resulting trust behavior toward the two trustees. Individual difference analyses showed a relationship between BLA and the CeA activation in the *preparation* phase and subjective trustworthiness and behavioral trust measures. More specifically, we first used a median split of *Δ investment* (**FIGURE 1B**) to distinguish learners from non-learners (i.e., those who adjusted their investment behavior less to the trustworthiness of the trustee), and then assessed how they differed in their amygdala activations. A Mann-Whitney *U*-test showed that during the *preparation phase*, the activation difference between untrustworthy–trustworthy trustee was significantly larger in learners than in non-learners in the BLA (*p*=0.0020, *u*=275.0, **FIGURE 3A**) and in the CeA (*p*=0.0336, *u*=350.0, **FIGURE 3D**). Moreover, the BLA activation differences between untrustworthy vs. trustworthy trustee in this phase correlated positively with behavioral trust (*Δ investment), r*_*s*_=+0.28, *p*=0.0255 (**FIGURE 3B**) and subjective trustworthiness (*Δ trustworthiness), r*_*s*_=+0.38, *p*=0.0026 (**FIGURE 3C**). CeA activation differences correlated with subjective trustworthiness ratings, *Δ trustworthiness, r*_*s*_=+0.31, p=0.0138 (**FIGURE 3F**), but not with behavioral trust, *Δ investment* (**FIGURE 3E**). While considering whether to trust or distrust a trustee, the CeA in learners thus seems primarily linked to evaluations of trustworthiness, whereas the BLA is additionally relevant for the actual behavioral outcome as well as whether someone efficiently learns to adapt behavior to the actually reciprocated trust or not. Moreover, these relationships are driven by stronger engagement for rounds with the untrustworthy (compared to the trustworthy) trustee, suggesting that what is coded is rather the absence than the presence of trust.

The neural responses in the *preparation* phase mainly provide insights into how the acquired information about a trustee’s trustworthiness drives the decisions of participants. The activation in the *outcome* evaluation phase, on the other hand, tells us about how this information is acquired and possibly updated. As outlined above, we observed overall activation in the BLA during *outcome* evaluation phase (**FIGURE 2**), and this may be linked to reward processing (Lüthi A and Lüscher C, 2014). Surprisingly, though, we did not find differences between the trustworthy and untrustworthy trustee in the BLA or CeA in the outcome phase, and neither did we find correlations with trust behavior and trustworthiness rating. We thus extended our analyses to subcortical regions with particularly strong anatomical and functional connections to the amygdala. These were the nucleus accumbens (NAc), as well as the dopaminergic midbrain, comprising substantia nigra and the ventral tegmental area (SN/VTA), relevant for encoding reward, and the bed nucleus of the stria terminalis (BST), relevant for encoding threat (Avery SN et al., 2016;Clauss JA et al., 2019;Siminski N et al., 2020).

When the *outcome* of th etrustee decision was presented, higher activation in the NAc, SN/VTA, and BST were observed for the trustworthy compared to the untrustworthy trustee (NAc *t=*+7.21, *p*<0.0001, SN/VTA: *t*=+3.31, *p*<0.0010, and BST *t*=+4.38, *p*<0.0001) (**FIGURE 4**). Moreover, the gain or loss (i.e., *back-transfer - investment amount*) correlated with NAc (*r*_*s*_=+0.19, *p*<0.0001) and BST (*r*_*s*_=+0.10, *p*<0.0001), but this was irrespective of the activation difference between trustworthy and untrustworthy trustee.

## DISCUSSION

Our previous study in BLA-damaged participants highlighted that the BLA is indispensable for learning to differentiate between trustworthy and untrustworthy trustees in the trust game (Rosenberger LA,Eisenegger C,Naef M,Terburg D,Fourie J,Stein DJ and van Honk J, 2019). This has important implications for our understanding of social decision-making in humans and, most likely, other mammals (O’Connell LA and Hofmann HA, 2012).

However, extending these findings to the neural networks connected to the amygdala in healthy, neurotypical, human participants is of the essence. Here we confirm the relevance of the BLA for distinguishing between trustworthy and untrustworthy trustees based on previous experience and how, in conjunction with the CeA, it plays a role in the guiding of trust behavior. Specifically, BLA activity was increased during the processing of the outcome of the trustee’s behavior but unselectively for trustworthy vs. untrustworthy trustee. Instead, we found increased activation in the NAc, BST, and SN/VTA for the trustworthy vs. untrustworthy trustee during outcome processing. Importantly, here we did not observe an activation difference between learners and non-learners. This could indicate that learners and non-learners processed the outcome in a similar fashion, suggesting that their understanding of the task and motivation were comparable. This further highlights the central role of the BLA for trust learning.

Indeed, we found the BLA to be most active during outcome evaluation, i.e., when participants learned whether their trust was reciprocated or not, suggesting that it plays an important role in acquiring beliefs about the trustworthiness of others. It appears, however, that the BLA is not directly involved in building specific outcome expectations during the *waiting* and *evaluation* phase. The BOLD response in the BLA was not modulated by the trustworthiness or the trustees’ back-transfer amount, unlike activity in the NAc, SN, and BST. This highlights that the BLA, although indispensable for learning whom to trust (Rosenberger LA,Eisenegger C,Naef M,Terburg D,Fourie J,Stein DJ and van Honk J, 2019), as indicated by our previous research, is only a component of a complex brain network for reward processing and social evaluation.

In addition, we found that while participants prepared for their next investment, the BLA together with the CeA exhibits increased activation for the untrustworthy trustee.

Importantly, this activation difference was only found in those participants who learned to differentiate between the trustees, indicating its role in (1) *guiding trust behavior* as BLA activation differences directly precede the participant’s investment behavior and also (2) in *trustworthiness evaluation*, as BLA and CeA BOLD responses correlated with the subjective rating after the experiment.

Nowadays, it is a well-established finding that a sub-population of BLA’s neurons selectively responds to reward, whereas other sub-populations either only respond to aversive stimuli (Pryce CR, 2018), or selectively increase their firing rate when the rewarding or aversive stimulus was unexpected, i.e., not predicted (Belova MA et al., 2007) (which means that something novel has to be learned about the environment). In the context of our findings, this view supports the notion that the BLA is relevant for encoding both the rewarding behavior of the trustworthy trustee and the aversive behavior of the untrustworthy trustee. Additionally, we can speculate that optimal performance in the trust game does not only rely on reward learning and threat detection, but also on predicting affective consequences based on abstract information. Supporting evidence for this theory can be found in a recent study in a patient with acquired complete bilateral amygdala lesions (patient SM, 49 years old, female), who showed impairments in making good predictions about what kind of written statements will induce fear (Cardinale EM et al., 2021).

The fact that we did not observe any habituation in any of the amygdala subregions (**SUPPLEMENTARY FIGURE 2**) indicates that the BLA not only responds to novel stimuli but is relevant for the continuous encoding and updating of information of social experiences. In the light of the recent debate on amygdala BOLD signal habituation (Geissberger N et al., 2020;Infantolino ZP et al., 2018;McDermott TJ et al., 2020;Plichta MM et al., 2012;Sladky R et al., 2012) this finding could be important for the development of additional tasks that robustly activate the amygdala.

While BLA’s activation during outcome evaluation suggests its involvement in discriminating and tracking outcome-specific effects, the CeA is involved in general motivational aspects of reward-related events (Corbit LH and Balleine BW, 2005) and, thus, might not play a role in the actual learning process in the *outcome* phase. Instead, we found it active during the *preparation* phase, which immediately preceded the *investment* phase. This could indicate that the CeA is regulated by the BLA output, which has been demonstrated before for a different task in a cross-species model (Terburg D et al., 2018). As CeA activity was increased before the participant’s investment, it might play a role in controlling trust behavior. More importantly, CeA activity during the preparation phase correlated with the subjective rating of trustworthiness of the trustee, indicating that it could be relevant for encoding the affective value attached to the trustee.

During *outcome* evaluation, we observed increased activation in the bed nucleus of the stria terminalis (BST), which, together with the CeA, is considered the *extended amygdala* complex (Alheid G and Heimer L, 1988;de Olmos JS and Heimer L, 1999). The BST has been suggested to play a role in both reward processing and social cognition (O’Connell LA and Hofmann HA, 2011) and exhibits strong connections to the NAc (Avery SN et al., 2014). While the CeA is associated with fast fear responses (e.g., startle reflex), the BST is responsible for slower affective learning processes (Gewirtz JC et al., 1998) and has been linked to adaptive and maladaptive responses to sustained stress and threat (Avery SN,Clauss JA and Blackford JU, 2016;Somerville LH et al., 2013). Of note, the BST plays a particular role in dealing with unpredictable threat (Goode TD et al., 2019), which could be the case in an uncertain social investment. However, these two views are still part of ongoing debates (Pedersen WS et al., 2019;Shackman AJ and Fox AS, 2016). Most recently, the BST was shown to be more involved in fear-related anticipation processes, whereas the CeA was linked to threat confrontation (Siminski N,Böhme S,Zeller J,Becker M,Bruchmann M,Hofmann D,Breuer F,Mühlberger A,Schiele M and Weber H, 2020). In this study we found the BST to be involved in the *outcome* evaluation phase. Based on the literature, it could be expected that the BST would show more activation for the aversive *untrustworthy* trustee, which was not the case. Instead, we observed that the BOLD responses of BST and NAc were both more activated by the trustworthy trustee. The NAc and other striatal areas are known to be involved in evaluating the trustees trustworthiness based on their back-transfer behavior (Baumgartner T et al., 2008;Delgado MR et al., 2005;King-Casas B et al., 2005) and amygdala to NAc coactivation is relevant for social decision making (Haruno M et al., 2014). Rodent research has shown that BLA to NAc connections mediate reward learning (Namburi P et al., 2015;Sesack SR and Grace AA, 2010). Importantly, stimulus-evoked excitation of NAc neurons depends on input from the BLA and is required for dopamine to enhance the stimulus-evoked firing of NAc neurons, ultimately, leading to reward-seeking behavior (Ambroggi F et al., 2008). This could mean that both regions might engage in a synergetic fashion, where the NAc would be particularly relevant for tracking rewards. The BST, on the other hand, could be responsible for increasing arousal as generous investments in the trustworthy trustee also entail a potential threat of betrayal. These findings suggest a functional dissociation between reward and risk evaluation based on the observed outcome of one’s behavior, which appeared to be comparable in non-learners, and the mechanisms of trust learning.

In sum, we confirm that the BLA is indeed involved in learning whom to trust and that observations from amygdala-lesioned participants can be translated to healthy neurotypical participants. Additionally, our fine-grained, time-resolved analyses of the amygdala subnuclei and the functionally-connected brain areas provide important insights into different cognitive mechanisms involved in trust learning. We found that the BLA is relevant for *discriminating* between trustworthy and untrustworthy trustees based on previous experience and for *optimizing trust behavior*. Only in those participants who learned to optimize their investments, we found selectively more activation in the BLA during the planning of a new investment that required trust. The BLA was also active during outcome evaluation suggesting its involvement in the process of *belief formation* based on the trustees’ back-transfer amount. As we did not observe a difference between the trustworthy or untrustworthy trustee, we can assume that encoding of potential rewards and risks is mediated by the NAc and BST, respectively, which showed a selectively increased activity for the trustworthy trustee or an increased investment. Finally, the CeA is known to receive inputs from the BLA and BST, and exhibited the largest BOLD response during the *planning* phase. CeA activity did not correlate with the participant’s trust behavior, however, there was a correlation with the participant’s subjective belief of the trustees’ trustworthiness. This suggests that the CeA could encode subjective value, possibly also indirectly affecting trust behavior via the BLA. Taken together, our work suggests that there is a high demand for translational work on the amygdala, its subnuclei, and connected brain regions. Based on the present results, we propose that careful variations of the trust game in combination with computational modeling may serve as an experimental model to further uncover the neural mechanisms underlying human social cognition.

## ACKNOWLEDGMENTS

We thank Helena Hartmann for her help in collecting the data. This work was partially funded by a grant awarded to Claus Lamm from the Austrian Science Fund (FWF P29150). Claus Lamm and Lisa Rosenberger acknowledge funding from the Vienna Science and Technology Fund (WWTF VRG13-007).

## SUPPLEMENTARY INFORMATION

Supplemental Information can be found online at [tbc].

## AUTHOR CONTRIBUTIONS

Conceptualization and Methodology, R.S., F.R., L.R., J.v.H., C.L.; Investigation, F.R.; Formal Analysis, R.S., F.R.; Writing – Original Draft, R.S., F.R., L.R., J.v.H., C.L.; Writing – Review & Editing, R.S., F.R., L.R., J.v.H., C.L.; Funding Acquisition, C.L.

## DECLARATION OF INTERESTS

The authors declare no competing interests.

## SUPPLEMENT

**Supplementary Figure S1.**
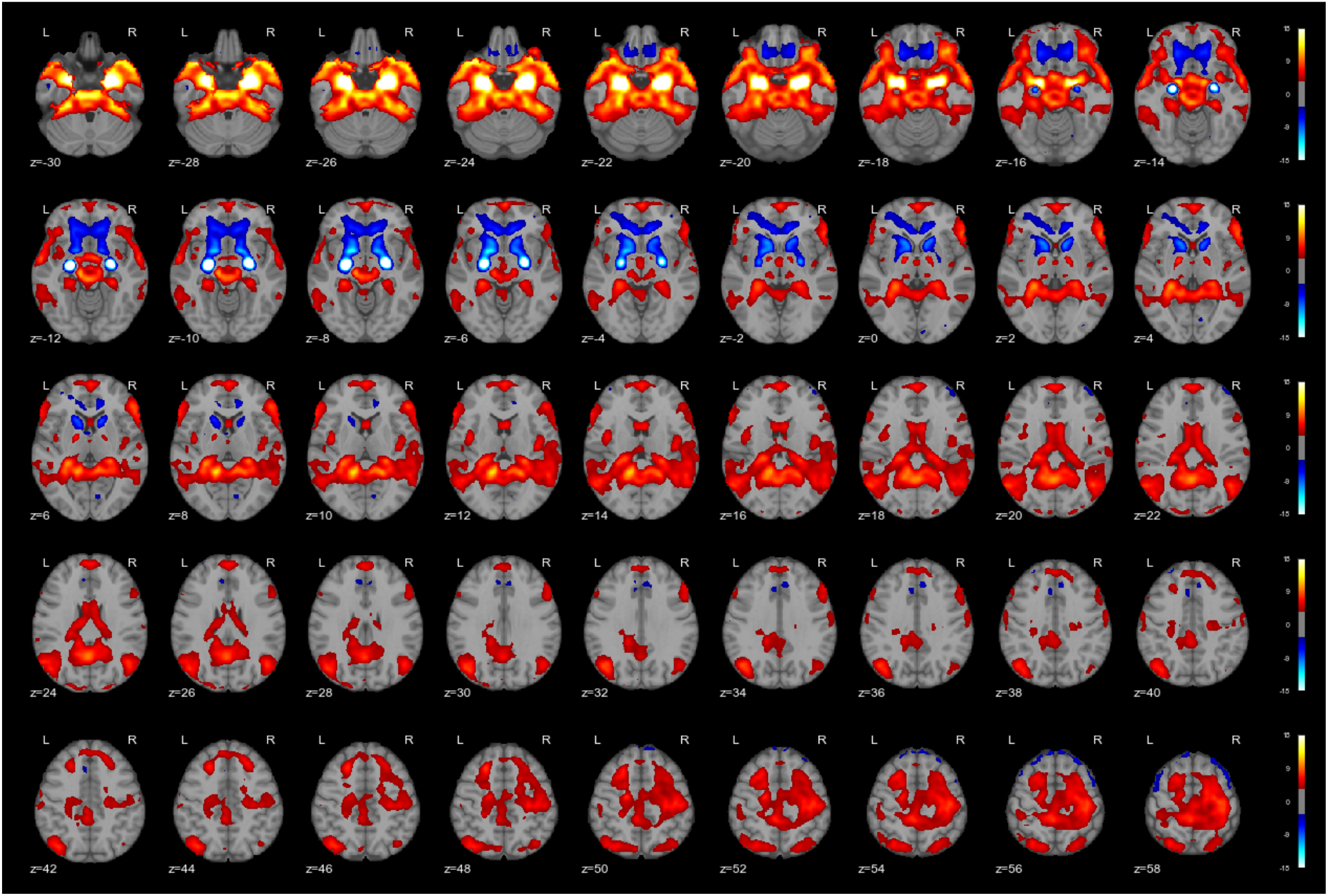
Differences in functional connectivity of BLA>CeA (hot) and CeA>BLA (cool).

**Supplementary Figure S2.**
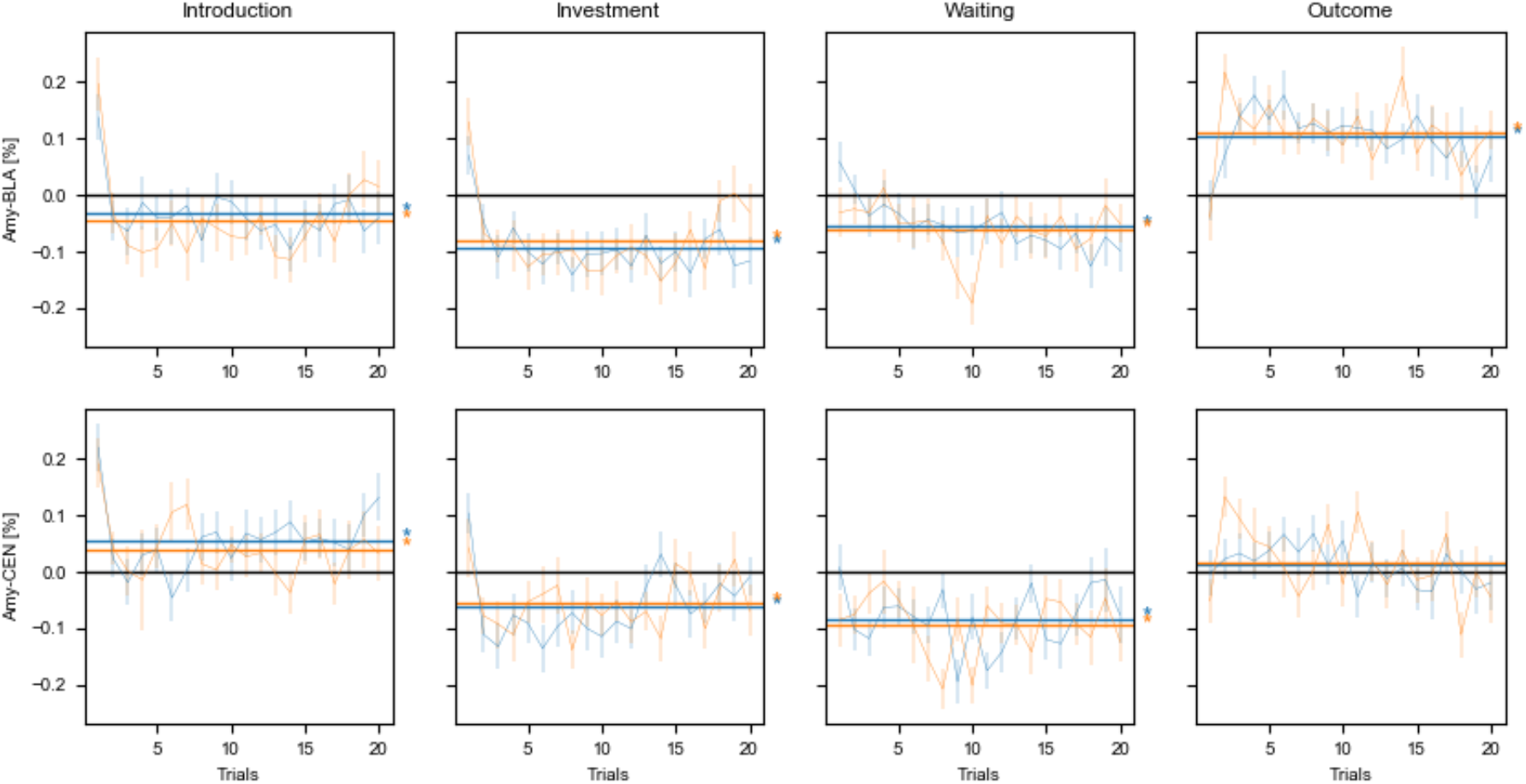
No evidence for amygdala habituation. Averaged percent signal change for the different task phases for the trustworthy (blue) and untrustworthy trustee (orange).

## Notes

### Competing Interest Statement

The authors have declared no competing interest.

